# Reversed sex-biased mutation rates for indels and base substitutions in *Drosophila melanogaster*

**DOI:** 10.1101/2020.04.08.031336

**Authors:** Lauri Törmä, Claire Burny, Christian Schlötterer

**Affiliations:** Institut für Populationsgenetik, Vetmeduni Vienna, Vienna, Austria; Vienna Graduate School of Population Genetics, Vetmeduni Vienna, Vienna, Austria

**Keywords:** Sex-biased mutations, *spellchecker1*, faster-X evolution, *Drosophila melanogaster*

## Abstract

Sex biases in mutation rates may affect the rate of adaptive evolution. In many species, males have higher mutation rates than females when single nucleotide variants (SNVs) are considered. In contrast, indel mutations in humans and chimpanzees are female-biased. In *Drosophila melanogaster*, direct estimates of mutation rates did not uncover sex differences, but a recent analysis suggested the presence of male-biased SNVs mutations. Here we study the sex-specific mutation processes using mutation accumulation data from mismatch-repair deficient *D. melanogaster*. We find that sex differences in flies are similar to the ones observed in humans: a higher mutation rate for SNVs in males and a higher indel rate in females. These results have major implications for the study of neutral variation and adaptation in *Drosophila*.

## Introduction

Mutations are the ultimate source of novelty in evolution. However, mutations do not occur at a constant rate across the genome (1, 2), between taxa (3), or sexes (4). The neutral substitution rate is determined by the primary mutation rate (5) and it needs to be considered when studying the intensity of selection from divergence data. If mutation rates differ between sexes, genes are expected to evolve at different rates depending on which chromosome they are located (6). For example, a higher mutation rate in males would result in a lower substitution rate on the X chromosome compared to the autosomes, while the Y chromosome has the highest substitution rate.

Higher male mutation rates have been observed for SNVs in mammals (7–9), birds (10, 11), salmonid fishes (12) and plants (13, 14). For indels, the observed sex-biases show no consistent trend across species. Human and chimpanzee females have a higher indel rate than males (15–17). In barn swallows, a sex-specific effect was shown for the mutation rate of a single microsatellite locus (18, 19) while in mice and rats indels are male-biased (20).

While sex-specific mutation rates are frequently observed, the underlying processes are highly taxon and mutation-type specific. In mammals, the higher rate of SNV mutations in males has been explained by more germline cell divisions in testes (9, 21). This explanation was challenged by the observation that even in young parents which have a similar number of germline cell divisions, males have a three times higher mutation rate than females (22). Moreover, the male-to-female mutation rate ratio barely increases with age, which is in conflict with the explanation involving germline divisions (22). In plants, explanations for sex-specific mutation rates include differences in per-replication mutation rates between the sex chromosomes (13), as well as different contributions of somatic mutations in males and females (14). For indels, the quiescent state of the oocyte may be responsible for the higher female mutation rate in humans (17). Similar to yeast (23), the quiescent oocyte is expected to acquire mutations not during replication, but from other sources, such as double-stranded breaks (17). In mice and rats, the male-bias of indel mutation rates is consistent with the hypothesis that DNA replication errors are the major source of mutations (20).

Consistent with a similar number of cell divisions for male and female germlines - on average 35.5 in males and 34.5 in females (24) - early sequencing studies in *Drosophila* found similar mutation rates for both sexes (25–29). Whole-genome comparisons with other *Drosophila* species provided conflicting results. These studies reported higher (30), equal (31), and lower (32) neutral substitution rates for the X chromosome. A recent analysis comparing *D. melanogaster*, *D. simulans*, and *D. yakuba* genomes challenged some of these findings and suggested that the X to autosomal (X/A) ratio of mutation rates could range from 0.81 to 0.93, depending on the type of neutral site used for the estimation (33). Further evidence for male-biased base substitution rates comes from the comparison of neo-Y and neo-X chromosomes in *D. miranda*, which showed higher substitution rates on the neo-Y chromosome (34). Since polymorphism and divergence patterns are strongly affected by selection, it is particularly noteworthy that a trend for male-biased mutations — although nonsignificant — was seen in mutation accumulation lines (35).

In this study, we estimate *de novo* mutation rates on the X chromosome and the autosomes using data from a mutation accumulation study mismatch-repair (MMR) deficient *D. melanogaster* strain consisting of 7,345 new single nucleotide variants (SNV) and 5,672 indels (36). We find opposing patterns of sex-specific mutation rates for SNVs and indels. While males have a higher mutation rate for SNVs (X/A mutation rates ratio: 0.876), indels are more common in females (X/A mutation rates ratio: 1.959). We show that wild-type females also have a higher indel mutation rate and conclude that our results are not an artifact from the mismatch repair deficiency, but reflect genuine sex-specific differences in mutation processes. These results have major implications for the interpretation of neutral variation and adaptation in *D. melanogaster*.

## Results and discussion

We scrutinized sex-specific mutation rates in *D. melanogaster* by comparing the *de novo* mutations on the X chromosome and the autosomes in a mismatch repair deficient genetic background, lacking a functional copy of the MutS homolog *spellchecker1* (37). Because the X chromosome spends more time in females than in males, a higher mutation rate in males leads to a higher mutation rate on the autosomes compared to the X chromosome. Consistent with previous results (33), we found that the X/A ratio of mutation rates was 0.876 (Poisson test, 95% Confidence Interval (CI): 0.823-0.933, p-value=2.923×10^−5^) resulting from a X chromosomal mutation rate of 8.71×10^−7^ (95% Poisson CI: 8.23×10^−7^−9.23×10^−7^) and an autosomal rate of 9.94×10^−7^ (95% Poisson CI: 9.69×10^−7^−1.02×10^−6^) (Figure 1a & Table 1). The difference between chromosomes still persists after controlling for the base content between the chromosomes (Figure 1b & Table 1).

**Table 1.**
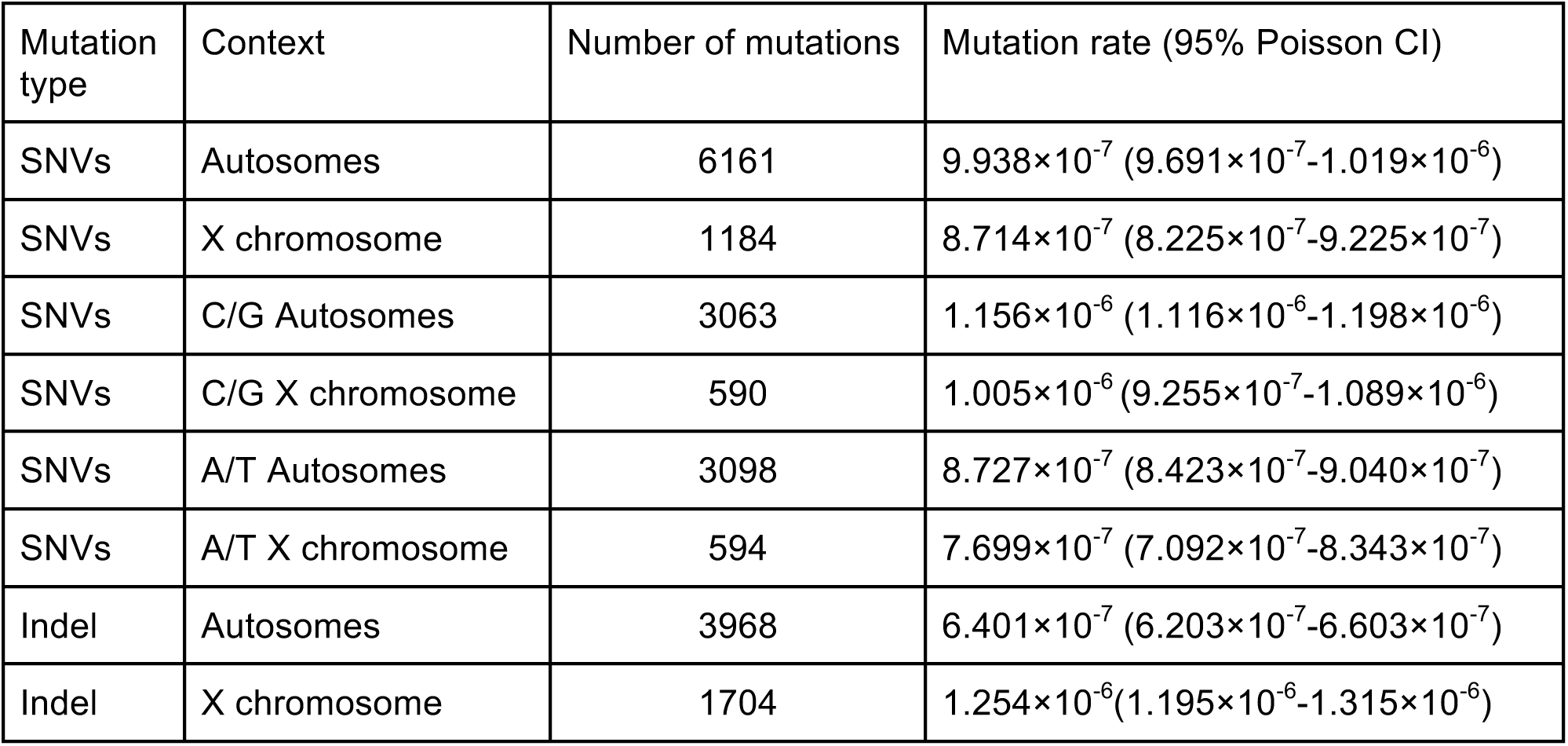
Mutation rate estimates.

| Mutation type | Context | Number of mutations | Mutation rate (95% Poisson CI) |
| --- | --- | --- | --- |
| SNVs | Autosomes | 6161 | $9.938 \times 10^{-7}$ ($9.691 \times 10^{-7}$-$1.019 \times 10^{-6}$) |
| SNVs | X chromosome | 1184 | $8.714 \times 10^{-7}$ ($8.225 \times 10^{-7}$-$9.225 \times 10^{-7}$) |
| SNVs | C/G Autosomes | 3063 | $1.156 \times 10^{-6}$ ($1.116 \times 10^{-6}$-$1.198 \times 10^{-6}$) |
| SNVs | C/G X chromosome | 590 | $1.005 \times 10^{-6}$ ($9.255 \times 10^{-7}$-$1.089 \times 10^{-6}$) |
| SNVs | A/T Autosomes | 3098 | $8.727 \times 10^{-7}$ ($8.423 \times 10^{-7}$-$9.040 \times 10^{-7}$) |
| SNVs | A/T X chromosome | 594 | $7.699 \times 10^{-7}$ ($7.092 \times 10^{-7}$-$8.343 \times 10^{-7}$) |
| Indel | Autosomes | 3968 | $6.401 \times 10^{-7}$ ($6.203 \times 10^{-7}$-$6.603 \times 10^{-7}$) |
| Indel | X chromosome | 1704 | $1.254 \times 10^{-6}$ ($1.195 \times 10^{-6}$-$1.315 \times 10^{-6}$) |

**Figure 1.**
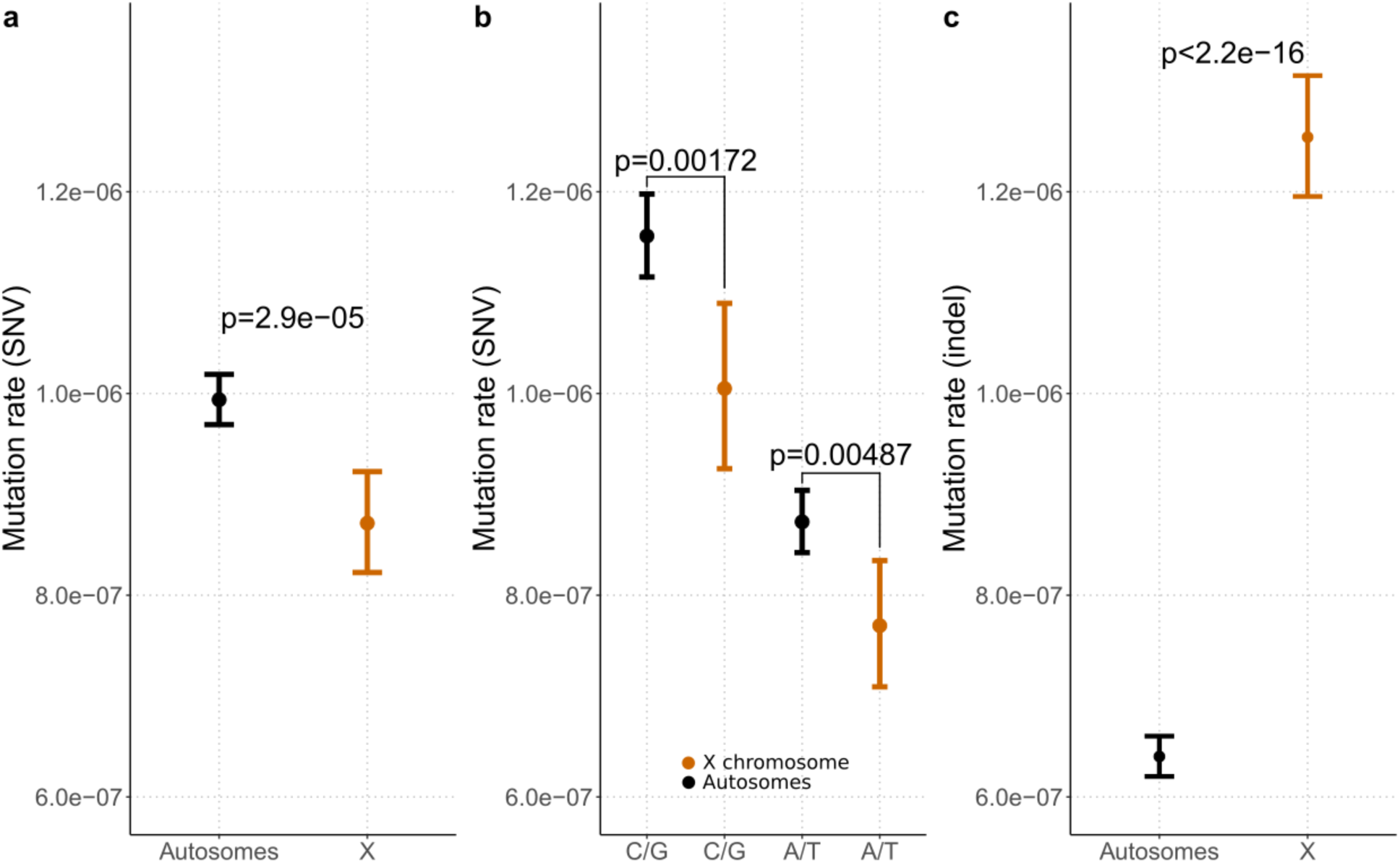
Mutation rate differences between the autosomes (black) and the X chromosome (orange). a) SNV mutation rates across all sites and b) separated for A/T and G/C pairs. c) Indel mutation rates across all sites. The bars represent 95% confidence intervals using exact Poisson tests, the dots represent the actual estimates.

With indel rates being sex-specific in multiple species (17–20), we tested for sex-specific indel rates in *Drosophila*. The indel mutation rate on the X chromosome was 1.26×10^−6^ (95% Poisson CI: 1.20×10^−6^−1.32×10^−6^), which is almost twice as high as the autosomal rate of 6.42×10^−7^ (95% Poisson CI: 6.22×10^−7^−6.62×10^−7^), resulting in a significant rate ratio of 1.959 (95% Poisson CI: 1.850−2.074, p-value<2.2×10^−16^) (Figure 1c & Table 1).

Indels occur more frequently on stretches of short tandem repeats (microsatellites) due to DNA replication slippage (38–40). Because microsatellite repeat number is an important factor in determining the indel mutation rate (41–43), the inferred sex-specific indel mutation rate may be the result of the heterogeneity in repeat length distribution between the X chromosome and the autosomes (44). We accounted for possible differences in the distribution of repeat length on the X and the autosomes by fitting a binomial generalized additive model (GAM, (45); see Methods) separately for mono- and di-nucleotide repeats. The indel mutation rate changes significantly with repeat length for both the homopolymers and dinucleotides (>5 degrees of freedom (dof), χ^2^= 5,046, p<2×10^−16^; >6 dof, χ2 = 1,676, p<2×10^−16^ respectively), but not in a monotonous way. The mutation rate increases for homopolymer runs up to 10-12 bp for (Figure 2a) and for dinucleotides up to 9-10 repeat units (Figure 2b). Microsatellites with more repeats were less likely to mutate and also less common. We found that the indel rate is significantly higher on the X chromosome for homopolymers (Odds Ratio X/A (OR)=1.163, 95% CI: 1.093-1.237, deviance analysis p=2.049×10^−6^) (Figure 2a) and dinucleotides (OR=1.291, 95% CI: 1.074-1.553, deviance analysis p=7.184×10^−3^) (Figure 2b). The highly consistent pattern across the two different repeat types strongly suggests that female flies have a higher indel mutation rate than males.

**Figure 2.**
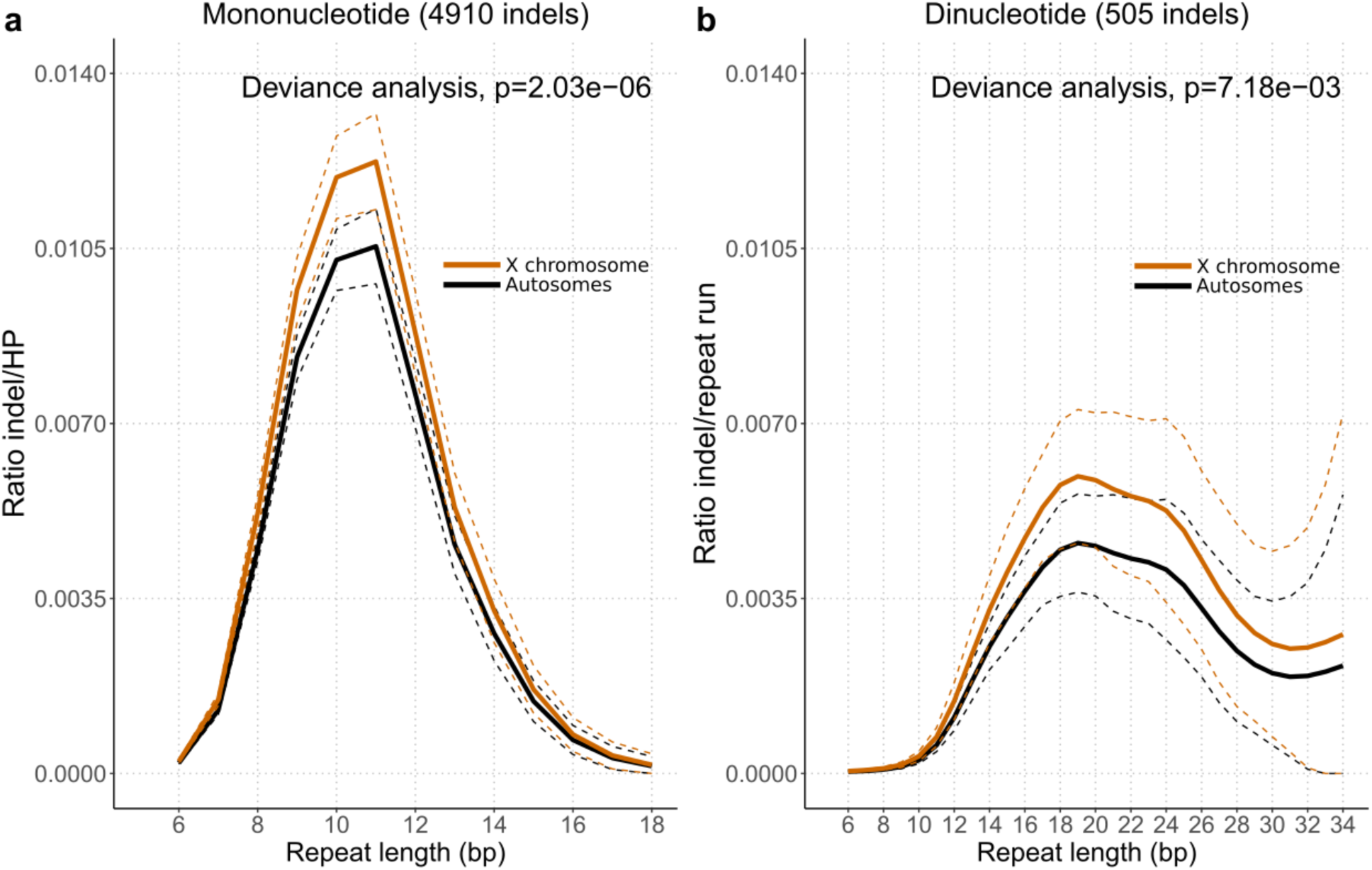
Occurence of indels in mononucleotide (a) and dinucleotide (b) repeats on the autosomes (black) and the X chromosome (orange). The x-axis represents the repeat length. The y-axis represents the ratio of indels normalized for the prevalence of repeat type in the genome, as described in (43). Binomial generalized additive models with cubic spline have been fitted. The predicted values are in plain lines, 95% confidence intervals are represented in dotted lines. The reported p-values are obtained from a deviance analysis contrasting a model with and without the chromosomal status (autosomes or X) covariate.

To rule out that the sex-specific difference in indel rates is an artifact of the mismatch repair system deficiency, we studied indel mutations in a natural *Drosophila* population. Using variant calls from the *Drosophila melanogaster* Genetic Reference Panel (DGRP) (46), we compared the number of indels to the number of SNVs between the X chromosome and the autosomes. We found an odds ratio significantly different from 1 (OR=1.320, 95% CI: 1.309-1.332, Fisher’s exact test, p<2.2×10^−16^), indicating that more indels occur on the X chromosome which confirms the female-biased indel mutation rate also in wild-type flies.

While we find the same sex-biased indel mutation rate as in humans and chimpanzees (17), we consider it unlikely that the interpretation of this sex-specific difference can be applied to *Drosophila*. For humans and chimpanzees, the quiescent state of the female oocyte has been attributed to the higher indel rate in females (17). This is however unlikely to cause the pronounced differences between the X chromosome and the autosomes in our experiment because it was designed such that young females were fertilized, leaving little opportunity for oocytes to enter a quiescent state. The mismatch repair targets replicating cells (47), which implies that most of the observed mutations did not occur in the quiescent phase, but during replication. Furthermore, the majority of indels (95.47% in mono- and di-nucleotides) occur on microsatellites arguing in favor of replication errors. Hence, we conclude that, at least in *Drosophila*, the higher indel rate in females cannot be explained by the quiescent phase of the oocytes, but rather other sex-specific differences in replication and/or DNA repair must be responsible for these differences.

One particularly interesting aspect of our study is that we detect sex-specific mutation rates for two types of mutations, SNVs and indels, but the sex-bias is reversed between them (Figures 1a, 1c). It has been proposed that the mutation rate has evolved as a balance between the cost in fitness due to accurate replication, repair, and deleterious mutations and the benefit of fitness increase due to advantageous mutations (48). In the case of sex-biased mutation rates, this implies that the cost/benefit balance differs between sexes (49). Since we show a different sex bias for SNVs and indels, it is difficult to reconcile a simple cost/benefit balance with our data. Rather, we consider it more likely that DNA damage and repair are sex-specific processes with their own signatures. The same conclusion was reached recently in humans (22).

The impact of the different sex bias for SNVs and indels is nicely illustrated by the different length distributions of AT-microsatellites on the X chromosome and autosomes in *D. melanogaster* (44). Microsatellite length may be explained by an equilibrium process between replication slippage generating longer repeats and base substitutions shortening the microsatellite (50). Hence, it was previously proposed that the difference in length distribution in *D. melanogaster* may be either explained by a higher slippage rate on the X chromosome or a higher base substitution rate in males (44). Our study now demonstrates that both processes occur and their joint effects probably explain the heterogeneous microsatellite length distribution between the X chromosome and autosomes.

In this study, we demonstrated that increasing the mutation rate in mutation accumulation lines is a powerful approach to study mutation processes, as it overcomes the typically encountered limitation of too few observations. We anticipate that the analysis of mutation accumulation lines with elevated mutation rates can provide a powerful method to study differences in the mutation process, either between sexes, as in this study, or between different genomic regions.

## Materials and methods

All statistical analyses were done with R (51) (version 3.5.0).

### Data and mutation rate estimations

We used mutations generated in (36) to perform a direct estimation of mutation rates. The VCF file for the DGRP Freeze 2.0 calls (46) was downloaded from: http://dgrp2.gnets.ncsu.edu/data.html in October 2019. We used the GATK tool CallableLoci (52) (version 4.0.12.0) to calculate the number of callable sites for each of the 7 lines from the same BAM files that were used in (36). We estimated the SNVs - using all SNVs and separated for A/T and G/C pairs - and indel mutation rates with the following formula: *m*/(*t* × *c*) where *m* is the total number of mutations across the 7 lines, *t* is equal to 10 generations, *c* is the total number of callable sites over each line. A 95% confidence interval was obtained by using an exact Poisson test with the poisson.test R function, as suggested in (53).

### Microsatellites and the model

We used a previously published R code (43) to search for homopolymer runs from BSgenome.Dmelanogaster.UCSC.dm6 (54). Dinucleotide repeats were annotated from the *D. melanogaster* reference genome release 6.24 using the microsatellite finder software MISA (55) (version 2.0). We included only microsatellites with i) at least 6 repeats and ii) a maximum length of 18 repeats (homopolymers) or 17 repeat units (dinucleotides), as longer microsatellites did not have any indel mutations. Following (43) we analyzed the occurrence of 5,415 (95.47%) indels separately for homopolymers and dinucleotides using binomial generalized additive models (45) with the non-linear effect of the repeat length. In addition we tested for differences between chromosomes by using the chromosomal status (autosomes or X) as covariate. The corresponding R function is mgcv::gam(cbind(#indels, #repeats - #indels) ~ Chr + s(Length, fx = FALSE, k=-1, bs = “cr”), family = binomial, data) (45, 56) (mgcv R package version 1.8-31), where *s* is a cubic spline, “#indels” is the number of indels per number of repeats, “#repeats” is the number of repeats of a given length and “Chr”, the chromosomal status. Two models, with (M1) and without (M0) the chromosome covariate “Chr”, were compared in a deviance analysis using the mgcv::anova.gam(M0, M1, test = “Chisq”) R function. Odds ratios (OR) and their 95% confidence intervals are obtained using the Miettinen-Nurminen method (57); OR=exp(β_*Chr*_ +/− −1.96 × *SE*), with β_*Chr*_ the slope associated with the chromosomal status (the autosomal status being the reference level) and *SE* its corresponding standard error. The p-values associated with the smoother of the length were reported from the summary R function. We reported the ratio of indels in both types of repeats normalized to the genome-wide number of repeats (Figure 2, y-axis).

### Code and data availability

The code (R and bash scripts) and intermediate files such as indel counts per repeat type will be accessible in the following github repository: ***, available upon publication.

## Author contribution

L. T., C. B. analyzed the data, L. T., C. B., C. S. wrote the paper, C.S. supervised the project.

## Acknowledgments

This work was supported by the Austrian Science Fund (FWF, grant W1225). We thank A. Futschik for statistical advice.

